# Chimeric LuxR Transcription Factors Rewire Natural Product Regulation

**DOI:** 10.1101/584987

**Authors:** Ruchira Mukherji, Somak Chowdhury, Pierre Stallforth

## Abstract

LuxR-type transcriptional activator proteins frequently flank bacterial biosynthetic gene clusters (BGCs) where they play a crucial role in regulating natural product formation. Only few bacterial BGCs are expressed under standard culturing conditions, thus modulation of flanking LuxRs is a powerful approach to activate silent clusters. Here, we show that exploiting the modular nature LuxR proteins and constructing chimeric LuxRs enables the activation of BGCs.

---

Ever since the beginning of the golden era of antibiotics, bacterial natural products have served as a key paradigm to fight infectious diseases^1,2^. However, in recent years, the number of multidrug-resistant pathogens has risen at an alarming rate. In this light, the search for novel anti-infective agents has become a matter of utmost priority. Genome mining has highlighted the untapped biosynthetic potential of bacterial genomes^3,4^. With plenty more natural products to be discovered, traditional screening approaches, however, rarely yield new chemical scaffolds^5^. Production of secondary metabolites is a great energy investment for bacteria, thus expression of biosynthetic genes is virtually always tightly regulated. In this report, we show that regulatory elements controlling bacterial biosynthetic gene clusters (BGCs) can be altered in order to modulate the conditions under which a particular secondary metabolite is produced. We focus on LuxR-type transcriptional activator proteins, which often flank bacterial BGCs^6^. These LuxR-family proteins serve as key regulatory elements and a deeper understanding of their effect on activation of adjacent BGCs allows determining conditions under which they can trigger natural product synthesis. LuxR-type regulators have been found to be a part of various quorum sensing (QS) systems that orchestrate BGC expression leading to the biosynthesis of multifarious natural products including siderophores, peptide antibiotics, polyketides, phenazines, pigments, and many others. LuxRs are response regulators that act as transcription factors upon binding a signal molecule produced by a cognate signal synthase. These LuxRs consist of two main domains: the N-terminal signal-binding domain (SBD) and C-terminal DNA binding domain (DBD). Upon binding to a cognate signal molecule (usually derivatives of acyl homoserine lactones, AHLs), the C-terminal part of the protein undergoes homodimerization and the dimer then acts as a functional transcriptional activator^7^. LuxRs specifically interact with discrete palindromic repeat motifs in the promoter region of their target genes, called lux boxes^8^. The modular nature of LuxRs and the genetic tractability of quorum sensing circuits makes them an ideal target for synthetic biology. Here, we describe the construction of chimeric LuxR proteins (LuxR^C^s) and demonstrate their functionality both in a synthetic system as well as in the native host (Fig. 1b, Supplementary Fig. S1). We hypothesize that a LuxR^C^ containing a well-characterized AHL-binding SBD and DBD of a LuxR associated with an unexpressed BGC, can jump-start the production of the corresponding metabolite. This strategy is a powerful addition to the natural product discovery toolbox.

**Fig. 1.**
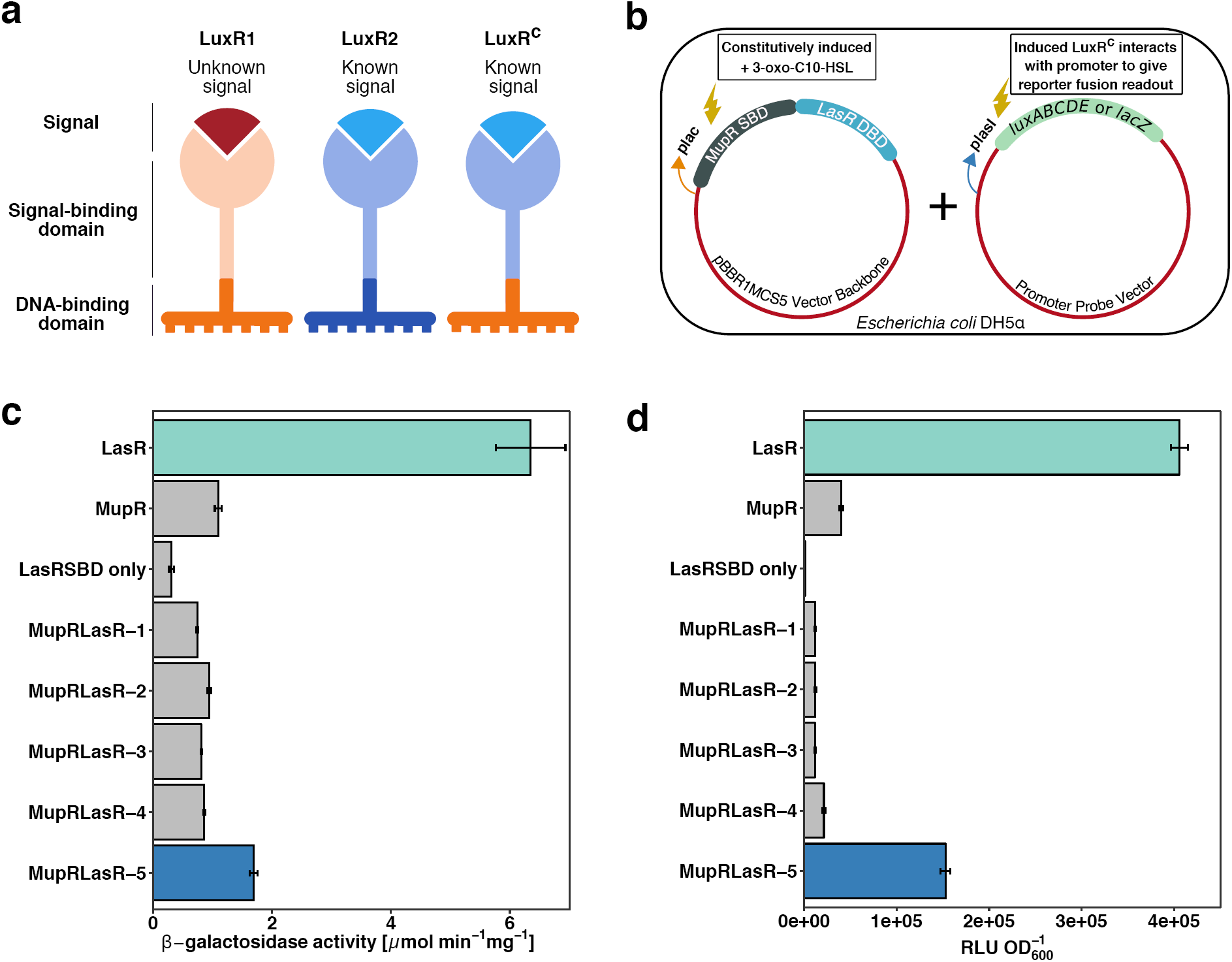
Schematic overview of the concept, the experimental set-up, and reporter gene fusion assays. **(a)** Architecture of LuxR chimera (LuxR^C^): chimeras are obtained by fusing the signal-binding domain (SBD) of LuxR2 with the DNA binding domain (DBD) of LuxR1. **(b)** Probing the functionality of LuxR^C^s in a synthetic system (*E. coli* DH5α): the activity of LuxR^C^ is determined by the quantification of β-galactosidase (*lacZ*) activity or luminescence (*luxABCDE*). **(c)** β-galactosidase activity obtained by the use of LuxR^C^_MupR-SBD::LasR-DBD_ variants as well as LuxRs. **(d)** Bioluminescence obtained by the use of LuxR^C^_MupR-SBD::LasR-DBD_ variants as well as LuxRs. This assay is more sensitive compared to β-galactosidase-based assay. LasR SBD served as negative control in both experiments.

As a proof of concept, we aimed at constructing a rewired synthetic system and test the biological functionality of LuxR^C^s in a heterologous host. We combined a SBD and a DBD from two different LuxRs into a single ORF. The MupR SBD was used, which is a part of QS regulatory machinery activating the production of the antibiotic mupirocin in *Pseudomonas* sp. QS1027^9^. This system responds to 3-oxo-C10 HSL. The DBD of LasR was used, which is a well-studied LuxR from *Pseudomonas aeruginosa*^10^. The vector encoding this LuxR^C^_MupR-SBD:: LasR-DBD_ was then co-transformed into *E. coli* DH5α along with a specific promoter probe vector (PPV) encoding the lux-box containing promoter region from *lasI*. This LasR-activated promoter controls the expression of *lacZ* or *luxABCDE*, resulting in β-galactosidase or luciferase production, respectively. A number of variants of LuxR^C^s, differing in the length of SBD or DBD, were tested and the results provided an insight into the modular nature of LuxRs. It was evident that the overall size of the *luxR*^*C*^s played an important role in generating a functional LuxR^C^. The size of *luxR*^C^ should match that of the native LuxR, which it aims to emulate; otherwise no transcriptional activity was observed (Fig. 1c,d). LuxR^C^s, which were able to dimerize and showed good transcriptional activity, comparable to the corresponding native LuxR, (Fig. 1) were then subjected to homology modeling. An interesting trend could be observed while analyzing the 3D-homology models of the LuxR^C^s compared to the native LuxR counterparts. 3D-structures of LuxR^C^ homodimers resembled that of the complete LuxR homodimer from which the AHL binding domain (SBD) was taken. Briefly, the homology model of functional LuxR^C^_LasR-SBD::MupR-DBD_ homodimer (Supplementary Fig. S2a) resembled the LasR homodimer (Supplementary Fig. S2b) and the functional LuxR^C^_MupR-SBD::LasR-DBD_ homodimer (Supplementary Fig. S2c) resembled the MupR homodimer (Supplementary Fig. S2d).

These preliminary studies regarding the generation of functional LuxR^C^s allowed us to design LuxR^C^s for *in-host* use. With the aim of rewiring the regulatory machinery of the mupirocin-producing BGC (*mup*) in *Pseudomonas* sp. QS1027, we used a LuxR^C^ with the LasR SBD from *Pseudomonas aeruginosa*, which binds to 3-oxo-C12 HSL, and the MupR DBD. In principle, this would allow us to change the signal that leads to mupirocin production from 3-oxo-C10 HSL to 3-oxo-C12 HSL. Thus, we introduced *luxR*^*C*^ variants episomally into *Pseudomonas* sp. QS1027 Δ*mupR*. Addition of the LasR signal, 3-oxo-C12 HSL should then lead to the expression of the *mup* BGC, which in turn leads to the production of mupirocin. Indeed, addition of 3-oxo-C12 HSL to *Pseudomonas* sp. QS1027Δ*mupR*, which produces LuxR^C^_LasR-SBD::MupR-DBD_ induced the biogenesis of mupirocin (Fig. 2a). Production titers were surprisingly high, almost matching that of wild type (WT) (Fig. 2a). This clearly showed that well-designed LuxR^C^ are functional and enable rewiring of natural product regulatory pathways. To further show the scope of our approach, we rewired the regulatory system for phenazine production in *Pseudomonas chlororaphis subsp. aurantiaca* (DSM19603). Production of orange-colored phenazines, specifically phenazine-1- carboxylic acid, required the LuxR, PhzR. Creation of Δ*phzR* mutant completely abolished phenazine production resulting in a strain with cream-colored colonies (Supplementary Fig. S4). Expression of a *luxR*^C^_LasR-SBD::PhzR-DBD_ in DSM19603 Δ*phzR* along with exogenous addition of 3-oxo-C12 HSL re-established the production of phenazine-1-carboxylic acid (Fig. 2b).

**Fig. 2.**
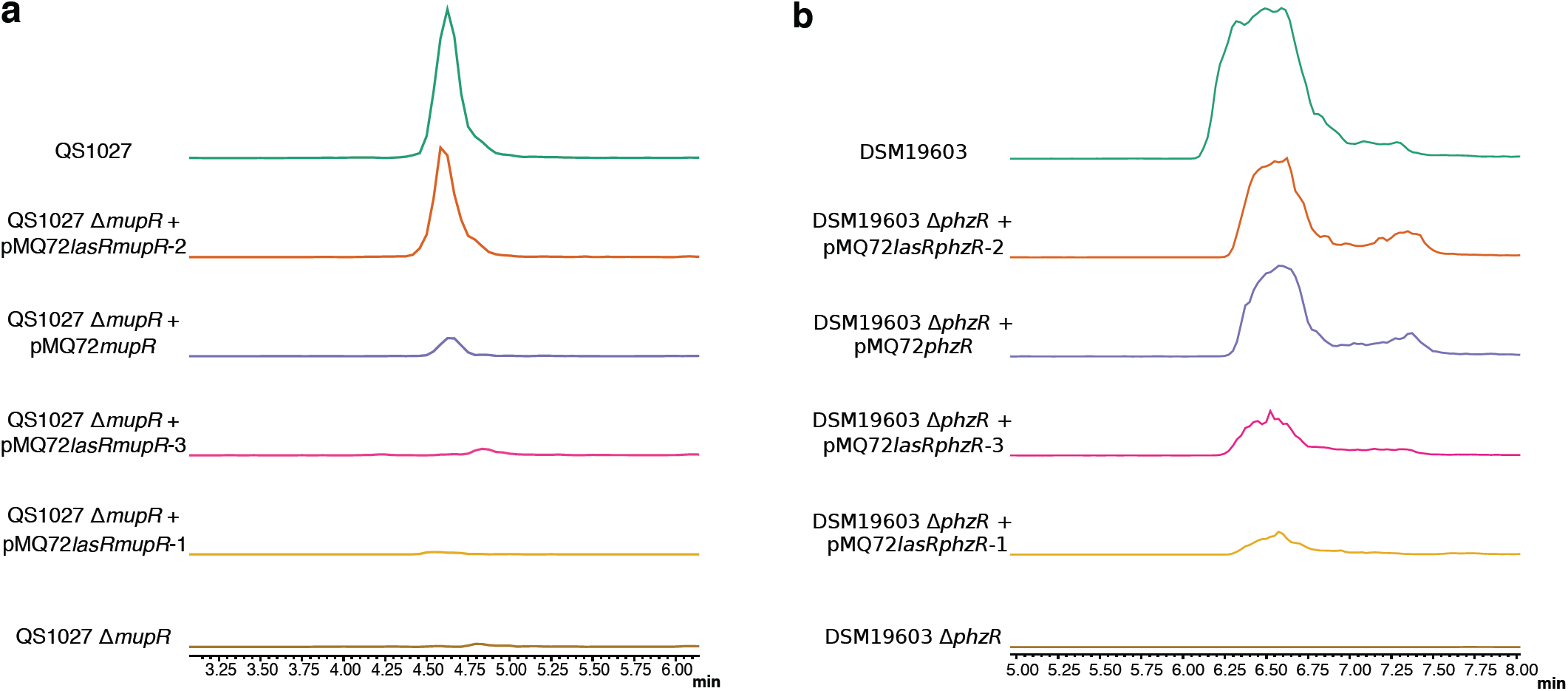
LC-MS-based analysis of natural product formation mediated by LuxR^C^ s. **(a)** Comparison of mupirocin formation amongst different mutants of *Pseudomonas* sp. QS1027 using single ion monitoring (SIM) at *m/z* = 501. Wild type (QS1027), (green) showed a high level of mupirocin synthesis and the null mutant Δ*mupR* did not produce any mupirocin, (brown). LuxR^C^_LasR-SBD::MupR-DBD_ (variant 2) was functional and restored mupirocin production in Δ*mupR* (orange). **(b)** Comparison of phenazine-1-carboxylic acid formation amongst different mutants of *P. chlororaphis* DSM19603 using SIM at *m/z* = 225. LuxR^C^_LasR- SBD::PhzR-DBD_ (variant 2) led to strong phenazine production in Δ*phzR* (orange).

Quorum sensing-controlled natural product production is ubiquitous in Gram-negative bacteria, in particular in *Pseudomonas* species^11^. Variants of these QS systems are known, where LuxRs have no cognate signal synthase – so-called orphan LuxRs or LuxR-solos^6,12^. LuxR-solo flanked BGCs are widespread yet often silent since the corresponding LuxR-signal is unknown^12^. Our approach allows activating these BGCs by the introduction of LuxR^C^s. Hence, new bioactive secondary metabolites can be accessed that would not be produced under standard laboratory conditions. Furthermore, our approach is particularly useful for synthetic biologists, who want to rewire the QS-mediated regulatory machineries and create artificial systems with precise control of signals that active specific BGCs.

## Methods

### Materials

*Pseudomonas sp.* QS1027 WT and Δ*mupR, Pseudomonas chlororaphis subsp. aurantiaca* (DSM19603) WT and Δ*phzR* were cultured in King’s Medium B (KB) and Nutrient Broth respectively at 28°C. *Escherichia coli DH5α* was grown in Luria Bertani (LB) medium at 37°C. Antibiotics, whenever required, were used in the following concentrations: ampicillin 100μg mL^-1^, gentamicin 15μg mL^-1^, chloramphenicol 50μg mL^-1^, and kanamycin 50μg mL^-1^. AHLs stocks were prepared in DMSO at a concentration of 1mg/mL and were added exogenously to the media, during the experiments, at a concentration of 1μM.

### Molecular methods, plasmid construction

Plasmids were constructed using the Gibson Assembly protocol (New England Biolabs). Primers were designed with http://nebuilder.neb.com. PCR reactions were performed using Q5® High-Fidelity 2x Master Mix (NEB) for cloning and sequencing otherwise DreamTaq Green PCR Master Mix 2x (Thermo Scientific, Darmstadt) was used. PCR amplicons were purified by agarose gel electrophoresis and extraction. Linearized doubly digested plasmids were assembled with appropriate PCR products using NEBuilder HiFi DNA Assembly Master Mix. All plasmids were sequenced prior to use.

For *E. coli* DH5α, a two-plasmid system was used to demonstrate the interaction of chimeric or complete LuxR with a specific promoter region. One plasmid contained *luxR*, and the other plasmid was a promoter probe vector (PPV), wherein a promoter of choice is cloned upstream to a reporter gene. Plasmid vector pBBR1MCS5 was used to clone *luxR* or the *luxR*^*C*^ downstream to the constitutive *E. coli* promoter *plac*. The pFU series of PPVs (a gift from the lab of Prof. Petra Dersch)^13^ was employed. pFU62 with *lacZ* as the reporter gene fusion or pFU36 with the *luxABCDE* cassette from *Photorhabdus luminescence* was used. *plasI* (*lasI* promoter, *P. aeruginosa* PA14) was amplified from the respective gDNA and subsequently cloned upstream to the reporter genes in the PPVs independently. Functional LuxR^C^ proteins were constructed from SBD of one *luxR* gene (approximately 480 bps) and DBD (approximately 230 bps) of another *luxR*^14^. SBD and DBD from two different *luxR*s were then assembled with linearized pBBR1MCS5 using NEB Hi-Fi DNA Assembly Master Mix. Plasmids were co-transformed into electro- competent *E. coli* DH5α with a specific PPV and tested in different reporter fusion assays for the ability of LuxR^C^ (compared to LuxRs), to interact with their respective promoter regions. LasR SBD alone served as negative control.

For *in-host* studies involving *Pseudomonas sp*. QS1027 Δ*mupR* and *P. chlororaphis* Δ*phzR*, a mobilizable expression vector (pMQ72) was used^15^. The vector encoded a LuxR or a LuxR^C^ under the control of the arabinose-inducible *pBAD* promoter. The vectors encoding the chimeras were called: pMQ72MupR, pMQ72LasRMupR, pMQ72PhzR and pMQ72LasRPhzR. A number of variants of pMQ72 encoding LuxR^C^s were constructed, differing mainly in the lengths of SBD or DBD of the candidate LuxRs.

### Synthetic and *in-host* experimental setups

*E. coli DH5α* cells were co-transformed and grown overnight in liquid medium (with ampicillin and gentamicin). The cultures were then diluted 1:100 and AHLs were added. The cultures were grown to late exponential phase and harvested for reporter gene fusion assays.

For *in-host* experiments, the pMQ72 vector backbone encoding either LuxRs or LuxR^C^s was first transformed into chemically competent *E. coli ET12567* pUZ8002, for conjugation, and grown on LB agar plates with kanamycin, chloramphenicol, and gentamicin. For biparental mating an overnight culture of the donor stain (*E. coli ET12567 pUZ8002* + plasmid) and acceptor strain (*Pseudomonas* sp. QS1027 Δ*mupR* or *P. chlororaphis* Δ*phzR*) were diluted 1:50 in their respective medium and grown till OD_600_ = 0.5. The two cultures were then mixed in different ratios 1:1, 2:1, 3:1 (donor: acceptor) and washed twice with deionized water. Mating mixtures were then spotted (25 µL) onto LB agar plate and incubated at 28 °C overnight. The mating spots were then suspended in LB medium (200 µL) and 100 µL of this suspension was then plated on LB agar plates with gentamicin and ampicillin. The plates were incubated at 28 °C for *ca.* 72 hours. Transconjugants were re-patched onto LB agar plates with gentamicin and ampicillin and incubated for 24 hours at 28 °C. Single colonies of *Pseudomonas* Δ*luxR* containing pMQ72*luxR*^*C*^ expression vector were used further for expression studies.

*Pseudomonas* sp. QS1027 Δ*mupR* and *P. chlororaphis* Δ*phzR* with different variants of LuxRs were grown overnight in medium with 0.4% glucose and gentamicin. The overnight cultures were pelleted and re-suspended in media with gentamicin, 3-oxo C12 HSL, and 100 mM arabinose to induce LuxR production. The culture was grown for 36 hours and extracted with ethyl acetate. The organic phase was dried over sodium sulfate and the solvent was removed *in vacuo*. LC-MS samples were prepared by dissolving the dried bacterial extracts in 200 µL methanol and filtration through PTFE filter membranes (0.2 µm). Samples were analyzed *via* LC-MS using positive mode single ion monitoring mode set at *m/z* 501 (mupirocin) and *m/z* 225 (phenazine-1-carboxylic acid).

### Reporter gene, β-galactosidase, bioluminescence assay

β-galactosidase activity was estimated in cell free extracts by measuring the intensity of yellow color obtained, from the breakdown of the substrate ONPG (420 nm). Enzyme activity and was calculated as follows: **OD_420_ × 6**,**75 × OD_600_^-1^ × (t_n_ – t_0_ [in min])^-1^ × Vol^-1^ [in mL]**, expressed as: μmol min^-1^ mg of protein^-1^. All assays were performed in biological triplicates. Reporter fusions emitting bioluminescence were measured in non-permeabilized cells with TECAN plate reader (InFinitePro-M200) using the luminescence mode. Results were expressed as relative light units (RLU)/OD_600_.

### LC-MS

analysis was performed on a Shimadzu System (LC-30AD, SPD-M30A, and LCMS-2020). The system is equipped with an electrospray ion source and a Kinetex® C18 column (50 × 2.1 mm, 1.7 µm, 100 Å, Phenomenex, for mupirocin) as well as a Kinetex Phenyl-hexyl column (50 × 2.1 mm, 1.7 µm, 100 Å, Phenomenex, for phenazine-1-carboxylic acid). Elution gradient: solvent A: H_2_O + 0.1% HCOOH, solvent B: acetonitrile + 0.1% HCOOH, gradient: 10% B for 0.5 min, 10% – 100% B in 8 min, 100% B for 3 min, flow rate: 0.7 mL/min, injection volume: 10 µL. LC-MS results were analyzed using LabSolutions Postrun and Browser (v.5.60).

## Author Contributions

PS and RM designed the experiments, RM conducted all experiments, PS, RM, and SC analyzed the data, PS and RM wrote the manuscript.

## Acknowledgements

We are grateful for financial support from the Leibniz Association. This work was supported by the collaborative research cluster ChemBioSys SFB1127 of the DFG and by an Exploration Grant of the Boehringer Ingelheim Foundation. We are grateful to the Europäischer Fonds für regionale Entwicklung (EFRE) for an instrument grant.

